# DNA Demethylation Is Dispensable for Venetoclax-HMA Synergy in Acute Myeloid Leukaemia

**DOI:** 10.64898/2026.04.13.718134

**Authors:** Aida Selimović-Pašić, Lisa Haglund, Maike Bensberg, Júlia Goldmann, Sandra Hellberg, Colm E. Nestor

## Abstract

The BCL-2 inhibitor venetoclax (VEN) in combination with hypomethylating agents (HMAs) has improved treatment responses in acute myeloid leukaemia (AML), but the mechanisms underlying their synergy remain unclear. We investigated the role of DNA demethylation in the enhanced cytotoxicity of VEN-HMA combinations. Using AML cell lines, we compared the effects of azacitidine (AZA), decitabine (DAC), cytarabine (ARA-C) and the DNMT1-selective inhibitor GSK-3685032 (GSK5032) with VEN. As expected, VEN showed strong synergy with AZA, DAC, and the DNA-damaging agent ARA-C, but not with GSK5032, despite the latter inducing extensive DNA demethylation. Genome-wide methylation profiling confirmed that loss of DNA methylation did not correlate with increased cytotoxicity or synergy with VEN. Moreover, combining GSK5032 with ARA-C did not enhance cytotoxicity, indicating that DNA demethylation and DNA damage do not act additively. Instead, synergy was consistently associated with the DNA damage-inducing properties of AZA, DAC, and ARA-C. Extensive DNA demethylation tended to antagonize VEN activity, suggesting that the epigenetic effects of HMAs may limit their synergistic potential. Overall, our findings demonstrate that DNA damage-related cytotoxicity, rather than DNA demethylation, is the dominant mechanism driving VEN–HMA synergy and provide evidence that VEN-mediated cytotoxicity arises primarily from genotoxic stress, supporting refinement of treatment strategies.

## Background

Hematological malignancies are characterized by abnormal cell survival driven by overexpression of anti-apoptotic proteins like BCL-2, promoting disease progression and therapeutic resistance [1]. In acute myeloid leukaemia (AML), the BCL-2 inhibitor Venetoclax (VEN) shows limited efficacy alone [2], but improves responses and survival when combined with hypomethylating agents (HMAs) such as 5-azacytidine (AZA) or decitabine (DAC) in elderly patients [3-5]. Nonetheless, toxicities and frequent relapses remain major challenges [6, 7], underscoring the need for a deeper understanding of the cellular and molecular dynamics of these combination therapies.

Preclinical studies suggest VEN and HMAs act synergistically, potentially driven by both epigenetic mechanisms [8-10] and non-epigenetic processes such as DNA damage [11]. However, the combined DNA demethylating effects and DNA damage response of conventional HMAs have made it challenging to determine which mechanism contributes to this synergy. We set out to clarify the relative contributions of DNA demethylation and DNA damage-related cytotoxicity to the observed synergy between VEN and HMAs, using the novel DNMT1-selective inhibitor GSK3685032 (GSK5032) [12]. With potent DNA demethylating activity and limited DNA damage-associated cytotoxicity [13], GSK5032 serves as a powerful tool to disentangle these two effects.

## Results

### Venetoclax exhibits synergy with HMAs, but not with DNMT1-selective GSK5032

To systematically dissect the mechanistic contributions of DNA demethylation and DNA damage to the VEN-HMA synergy, we used human AML cell lines as a well-established and reproducible model system (**Fig. 1A**). Initial drug-sensitivity assays were performed to evaluate the effects of AZA, DAC, GSK5032, and VEN as single agents in four AML cell lines (**Fig. 1B, Fig. S1A**) [14]. To separate DNA damage from demethylation effects, we also included ARA-C, a nucleoside analogue that induces DNA damage without demethylating activity. MOLM-13 and Kasumi-1 were more sensitive to VEN, while KG1-a and THP-1 appeared more resistant in line with previous studies [15-17] (**Fig. 1B, Fig. S1A-C**). GSK5032 displayed lower toxicity at both three and seven days compared to AZA and DAC, whereas ARA-C treatment induced the highest cytotoxicity in all cell lines (**Fig. 1B, Fig. S1C**).

**Fig. 1.**
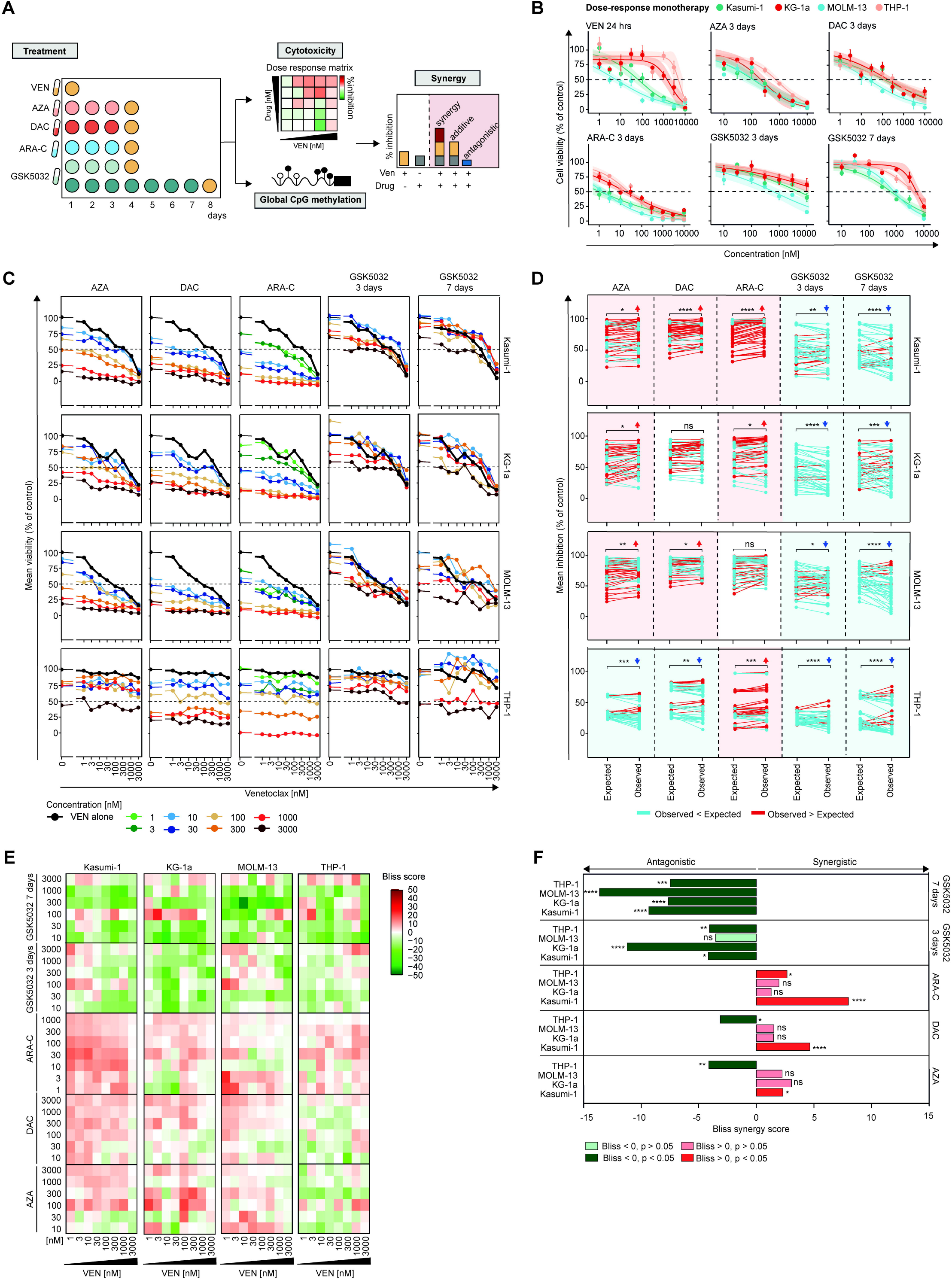
Lack of synergistic effect between a non-DNA-damaging DNA demethylating agent and Venetoclax in AML cell lines. **A** Overview of the experimental design where four AML cell lines (Kasumi-1, KG-1a, MOLM-13 and THP-1) were treated with increasing concentrations of 5-azacytidine (AZA), decitabine (DAC), GSK-3685032 (GSK5032) or Cytarabine (ARA-C) for three or seven days (only GSK5032), followed by 24-hrs Venetoclax (VEN) treatment. Treatment responses were assessed by (i) Alamar Blue assay for cell viability and (ii) whole-genome DNA methylation sequencing. The synergistic effects between VEN and each drug combination were evaluated using the Bliss model. **B** Monotherapy dose-response curves for all four AML cell lines following treatment with varying concentrations of AZA, DAC, ARA-C, GSK5032, or VEN for 24 hours, three days, or seven days (GSK5032 only). Cell viability was assessed by Alamar Blue assay. Mean ± SEM of triplicates are shown. **C** Mean viability expressed as percent relative to control of combination therapy with VEN and AZA, DAC, ARA-C, or GSK5032 after three and seven days (only for GSK5032). Black lines show effect of VEN treatment alone. **D** Comparison of observed combination effects to the expected effects calculated using the Bliss model, which assumes independent action between single agents. Lines connect each dose-combination pairing between the expected and observed effects. Red lines indicate combinations where the observed effect exceeds the expected effect (synergistic), while blue lines indicate combinations where it falls below (antagonistic). Statistical differences between the expected and observed effects were assessed using the Wilcoxon Rank Sum Test, where * = p < 0.05, ** = p < 0.01, *** = p < 0.001, and **** = p < 0.0001. **E** Synergy scores for treatments combining VEN with AZA, DAC, ARA-C or GSK5032 were calculated using the Bliss model. Heatmaps show the Bliss score for each combination, based on three replicates per combination. (F) The mean synergy score and the level of significance for each treatment combination; AZA-VEN, DAC-VEN, ARA-C-VEN and GSK5032-VEN divided per cell line. * = p < 0.05, ** = p < 0.01, *** = p < 0.001, and **** = p < 0.0001.

Since VEN has previously been shown to have synergistic effects with HMAs [8-11], we proceeded to determine whether similar synergistic effects could be observed using GSK5032. All AML cell lines were treated with varying concentrations of AZA, DAC, GSK5032, or ARA-C for three or seven days, followed by a 24-hour treatment with increasing concentrations of VEN, generating a comprehensive dose-response matrix of at least 63 different treatment combinations for synergy analysis. VEN combined with AZA, DAC, or ARA-C reduced cell viability compared to VEN alone across nearly all tested concentrations (**Fig. 1C**). Interestingly, while several dose combinations of VEN-GSK5032, particularly those containing higher concentrations of GSK5032, reduced cell viability compared to VEN monotherapy, lower concentrations of GSK5032 (10nM-100nM) did not lead to higher cytotoxicity. In contrast, AZA, DAC, and ARA-C reduced cell viability at even lower concentrations (**Fig. 1C**). Notably, the observed inhibition of VEN-GSK5032 was significantly lower than the expected effect, calculated using the Bliss Independence model, which assumes the two drugs act independently based on their monotherapy responses (**Fig. 1D**). In comparison, combinations with AZA, DAC, and ARA-C frequently resulted in significantly greater inhibition than expected for purely independent drug effects, suggesting potential synergy (**Fig. 1D**). Corresponding synergy scores confirmed that VEN-AZA and VEN-DAC combinations were more synergistic compared to VEN-GSK5032 (**Fig. 1E, Table S1**). Interestingly, VEN and GSK5032 more often exhibited antagonistic responses across all tested combinations (Bliss score < 0; **Fig. 1E, Table S1**) and cell lines, particularly after seven days of treatment with GSK5032 (**Fig. 1F, Fig. S2A-D, Table S2**). Since we observed no synergistic effect for GSK5032 but high synergy for ARA-C combined with VEN (**Fig. 1F**), we hypothesized that the observed synergy with AZA and DAC is driven by DNA damage rather than DNA demethylation.

### DNA demethylation does not contribute to Venetoclax synergy

Given the notably weaker or non-synergistic effects observed with GSK5032 compared to all other treatments, we proceeded to validate their DNA demethylating activity using our low-coverage whole methylome sequencing approach [13] (**Fig. 2A, Table S3-4**). Global DNA methylation levels obtained with this approach reflected methylation at long interspersed nucleotide elements (LINEs), which have long been used as a proxy for global methylation levels (**Fig. 2B, Table S5**) [18], validating the reliability of our approach. As expected, ARA-C did not affect DNA methylation levels (**Fig. 2C**), again indicating that synergistic effects observed with VEN are due to the DNA-damaging-properties of AZA, DAC and ARA-C. The greatest loss of DNA methylation was achieved with GSK5032, with as little as 15-26% of methylation remaining in Kasumi-1 and MOLM-13 after seven days treatment (**Fig. 2C**). Importantly, these cell lines also had the lowest Bliss synergy scores at seven days (Kasumi-1: mean Bliss score -9.32, p = 3.3e-07; MOLM-13: -13.62, p = 2.3e-11; **Fig. S2A, C**), clearly de-coupling loss of DNA methylation from the synergistic effects of VEN. Global methylation levels in Kasumi-1 and MOLM-13 cells significantly correlated with cell viability following treatment with AZA and GSK5032 (**Fig. 2D**). After seven days of GSK5032 treatment in Kasumi-1 cells, global methylation decreased to 26%, but cell viability remained high at 69%. In contrast, treatment with AZA for three days resulted in 66% global methylation, with only 15% cell viability (**Fig. 2D, Table S6**). Additionally, the two drugs showed markedly different synergistic effects in combination with VEN (**Fig. 1F**). GSK5032, which induced the strongest demethylation but had relatively limited cytotoxicity, showed minimal synergy with VEN. In contrast, AZA, which had more modest demethylating effects but greater cytotoxicity, was strongly synergistic with VEN. This suggests that DNA damage-related cytotoxic potency, rather than the extent of DNA demethylation, is the key driver of synergistic response of HMAs with VEN.

**Fig. 2.**
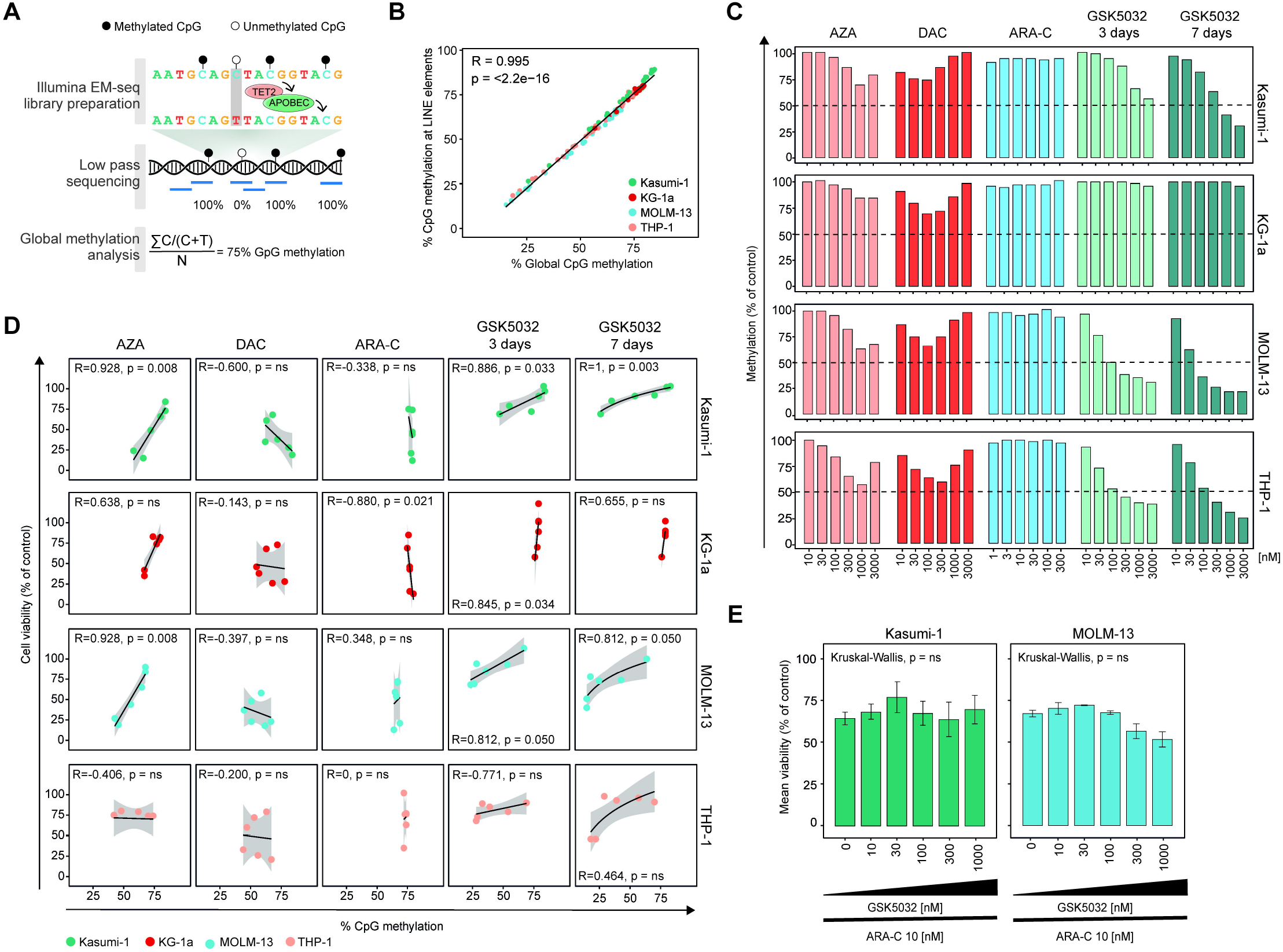
DNA demethylation does not enhance cytotoxicity. **A** Schematic overview of the enzymatic methyl-sequencing (EM-seq) workflow. EM-seq library preparation involves the oxidation of methylated and hydroxymethylated cytosines by TET2, followed by the deamination of unmethylated cytosines to uracil by APOBEC, which is subsequently read as thymine during sequencing. Libraries are sequenced at low coverage using paired-end sequencing. Global CpG methylation levels were calculated as ∑C / (C + T) / N, where C represents the number of methylated CpG sites, T the number of unmethylated CpG sites, and N the total number of CpG sites analyzed. **B** Correlation between average global CpG methylation assessed by low-coverage whole methylome sequencing and DNA methylation at long interspersed nucleotide elements (LINE) across all cell lines. Correlation was done with Spearman’s rank correlation. **C** Global CpG methylation relative to control in Kasumi-1, KG-1a, MOLM-13, and THP-1 cells after treatment with increasing concentrations of 5-azacytidine (AZA), decitabine (DAC), cytarabine (ARA-C), or GSK-3685032 (GSK5032) for three days or seven days (GSK5032 only). **D** Global CpG methylation and corresponding cell viability for each respective treatment and cell line. Generalized linear regression models were fitted to the data and shown in the plots. Spearman’s rank correlation coefficient was used to test for correlation between cell viability and DNA methylation. Grey band depicts the 95% confidence interval of the regression model. Correlation coefficients and corresponding p-values are shown in the figure. P-values greater than 0.05 are labelled as ‘ns’ (non-significant). **E** Mean viability after treatment with increasing concentrations of GSK5032 for three days and fixed concentration of ARA-C in Kasumi-1 and MOLM-13. Kruskal-Wallis was used to test for statistical differences.

### DNA damage, not DNA demethylation, drives synergistic cytotoxicity

To explore whether the DNA demethylating and DNA-damage-inducing properties of HMAs act together to induce toxicity, we combined low-dose GSK5032 with a fixed concentration of ARA-C. At these concentrations, GSK5032 induces robust DNA demethylation without affecting viability, enabling us to assess whether DNA demethylation is sufficient to drive strong combined cytotoxicity in VEN–HMA synergy. Cell viability was not significantly altered by the addition of increasing concentrations of GSK5032 compared to ARA-C alone (**Fig. 2E**). A slight trend toward decreased viability was observed in the MOLM-13 cell line, consistent with the modest cytotoxicity of GSK5032 monotherapy in these cells (**Fig. 1B**). Overall, these findings show that GSK5032 does not enhance ARA-C-mediated cytotoxicity, further suggesting that the DNA demethylating effects of HMAs contribute little to their toxicity in our model.

## Discussion

Venetoclax combined with hypomethylating has improved patient survival [3-5], but the mechanisms underlying their synergy remain unclear due to the dual effects of conventional HMAs on DNA demethylation and DNA damage. Using the DNMT1-selective inhibitor GSK5032, which induces robust DNA demethylation with limited cytotoxicity [13], we were able to dissect the specific contribution of DNA demethylation. Despite inducing global DNA demethylation, GSK5032 did not enhance VEN-mediated cytotoxicity, nor did it significantly increase cytotoxicity when combined with low-dose ARA-C. These findings suggest that the DNA-damaging properties of HMAs, rather than their epigenetic activity, are the primary drivers of the observed synergistic cytotoxicity with VEN.

Contrary to our findings, some studies have attributed the synergistic effects of VEN and HMAs to epigenetic regulation. For example, DNA demethylation has been shown to upregulate the pro-apoptotic protein PUMA [8], reactivate retroelements [9] and induce pyroptosis-related gene expression [10], driving synergy. Furthermore, previous studies have reported that HMAs can sensitize cells to VEN [19]. However, in our study, the VEN-resistant cell lines KG-1a and THP-1 were not sensitized by prior HMA exposure. In line with our findings, others suggest that synergy is mainly driven by non-epigenetic events such as upregulation of pro-apoptotic NOXA or BAX, driving mitochondrial membrane permeabilization [11, 20]. Although our experimental design cannot delineate the precise downstream molecular pathways involved, our results indicate that DNA demethylation alone is insufficient to drive synergistic VEN-mediated cytotoxicity, supporting a primary role for non-epigenetic mechanisms.

This *in vitro* study does not fully recapitulate the clinical treatment setting, where VEN is combined with AZA or DAC in 21–28-day cycles, with concurrent treatment for 5–7 days followed by VEN alone for the remainder of the cycle [5]. To specifically decouple the contribution of DNA methylation changes from other drug-induced effects, we selected drug concentrations and exposure time points that reliably produce measurable changes in global DNA demethylation prior to VEN exposure. Further validation in *in vivo* cancer models and primary cells will be essential to support clinical translation.

Taken together, by using GSK5032 we have been able, for the first time, to uncouple the demethylating effect from the cytotoxicity induced by conventional HMAs, revealing that DNA demethylation, unexpectedly, acts antagonistically rather than synergistically in combination with VEN. Our findings underscore the importance of understanding drug interactions in greater depth to optimize and refine combination therapies.

## Materials and Methods

### Cell lines

AML cell lines MOLM-13, Kasumi-1, KG-1a and THP-1 were ordered from German Collection of Microorganisms and Cell Cultures (DSMZ) and kept in humidified incubator at 37°C with 5% CO_2_. Cells were passaged every two or three days and routinely checked for mycoplasma contamination using the MycoAlert Mycoplasma Detection Kit (Lonza, LT07-318).

### Treatment with drugs

For monotherapy, cells received VEN for 24 and 48 hours; AZA, DAC, GSK5032, or ARA-C for three days; or GSK5032 for seven days. For combination treatment, cells were treated with AZA, DAC, GSK5032, or ARA-C for three days, followed by VEN for 24 hours (**Fig. 1A**). For the ARA-C/GSK5032 co-treatment, Kasumi-1 and MOLM-13 cells were exposed to a fixed concentration of ARA-C (10 nM) together with increasing concentrations of GSK5032 for three days. In all treatment settings, drugs were refreshed every 24 hours. Untreated controls were exposed to equivalent concentrations of DMSO or ultrapure H_2_O.

### Alamar Blue cell viability assay

Viability was analyzed with the Alamar Blue cell viability reagent (Thermo Fisher Scientific) and determined following the instructions provided by the manufacturer. Treatments were performed in triplicate. Dose response curves were generated in R (version 4.4.2) with the drc package version 3.0-1 fitting a three-parameter log-logistic function the fct = LL.3().

### Synergy analysis

The expected combination effect of VEN with AZA, DAC, ARA-C or GSK5032 was calculated using the Bliss Independence model as implemented in SynergyFinder Plus R package (version 3.14.0) [21]. Due to the nature of AlamarBlue analysis, where values from untreated control wells is subtracted from treated samples, calculated viability values can fall below 0% or exceed 100%, depending on the treatment effect. All values were therefore adjusted prior to analysis to constrain all values within the biologically relevant range of 0-100%; values below 0% were set to 0%, and those above 100% were set to 100%. Synergy scores of >0 was regarded as synergistic.

### DNA and RNA isolation

DNA and RNA were extracted using the Quick-DNA/RNA Kits (Zymo Research) according to the manufacturer’s instructions, with additional in-column RNase/DNase treatments.

DNA and RNA concentrations were measured using a NanoDrop 2000 spectrophotometer (Thermo Fisher Scientific).

### Quantitative PCR

200 ng RNA was reversed transcribed into cDNA using the High-Capacity cDNA Reverse Transcription Kit (Applied Biosystems), according to the manufacturer’s instructions. For each PCR reaction, 4.5 μl of diluted cDNA was combined with 5 μl of TaqMan™ Fast Universal PCR Master Mix (Thermo Fisher Scientific) and 0.5 μl of TaqMan probes (Thermo Fisher Scientific) for the following genes: *DAZL* (Hs00154706_m1), *GAGE12* (Hs04190947_gH), and *GAPDH* (Hs02758991_g1). qPCR was performed in technical duplicates on a QuantStudio™ 7 Flex Real-Time PCR System (Thermo Fisher Scientific) using the following thermal cycling conditions: 95°C for 20 seconds and 40 cycles of 95°C for 1 second and 60°C for 20 seconds. Expression of the known DNA methylation-sensitive genes *DAZL* and *GAGE12* was normalized to *GAPDH* expression. HMAs induced expression of both *DAZL* and *GAGE12* indicating local transcriptional changes to loss of DNA methylation (**Fig. S3**).

### Library preparation for enzymatic methyl-sequencing

Genomic DNA was sheared using the Covaris S220 ultrasonicator, yielding an average fragment size of 302 bp. Libraries were carried out using the NEBNext Enzymatic Methyl-seq Kit (New England Biolabs) following the manufacturer’s standard library protocol. Libraries were PCR-amplified for four cycles. The libraries were sequenced using low-coverage whole methylome sequencing with 150 bp paired-end reads, generating an average of 232,533 read pairs per sample (min 118,415 reads, max 453,764 reads). The sequencing was performed on MiSeq (Illumina) using the MiSeq Reagent Kit v2 (300 cycles) (Illumina). Only samples with CHG and CHH methylation levels < 1.5 % were included in downstream analysis to confirm successful library conversion.

### Global methylation analysis from low-coverage whole-genome enzymatic methyl-sequencing sequencing

Reads were trimmed with Trim Galore version 0.6.10 (https://github.com/FelixKrueger/TrimGalore) using default parameters including *–paired* and *--fastqc*. Sequences for pUC19 (M77789.2) and lambda (J02459.1) were added to the human hg38 reference genome (GCA_000001405.15_GRCh38_no_alt_analysis_set). Trimmed reads were aligned (bwameth.py) to the indexed genome (*bwameth*.*py index*) with bwa-meth version 0.2.7 (https://github.com/brentp/bwa-meth arXiv:1401.1129v2) with default parameters. Aligned reads were converted to bam format (*samtools view*) and sorted (*samtools sort*) using samtools version 1.13 [22]. Duplicate reads were marked with Picard *MarkDuplicates* (http://broadinstitute.github.io/picard) with default parameters and files were indexed with *samtools index*. DNA methylation was called with *MethylDackel extract* (version 0.6.1, https://github.com/dpryan79/MethylDackel) with trimmed reads by 5 bases from each end (*--nOT 6,5,6,5* and *--nOB 6,5,6,5*). Non-CpG methylation was extracted (*--CHG* and *--CHH*) to evaluate conversion. For CpG methylation calls, strand-specific information was merged (*MethylDackel extract –mergeContext --nOT 6,5,6,5 --nOB 6,5,6,5*). Additionally, the number of reads covering each CpG was extracted with *MethylDackel extract (--nOT 6,5,6,5; --nOB 6,5,6,5; --mergeContext, --counts*). From the strand-merged CpG methylation calls, global average CpG methylation levels were calculated after excluding both control DNAs (pUC19: M77789.2; lambda: J02459.1) and mitochondrial DNA (chrM) (**Table S3**).

### Data visualization

Data was visualized using ggplot2 (for line graphs, heatmaps, and bar graphs; version 3.5.1). Statistical analysis was conducted in R (version 4.4.2) as described if not stated otherwise.

## Code availability

All code used for data analysis is available at https://github.com/ColmNestorLab/Synergy.

## Acknowledgments

We acknowledge the Core Facility at the Faculty of Medicine and Health Sciences, Linköping University, for their support with equipment and assistance for sequencing. Sequencing was performed in collaboration and with support by Clinical Genomics Linköping, SciLifeLab, Sweden.

## Author Contributions

A.S.P., L.H., M.B., and J.G performed all the experiments; A.S.P., L.H., S.H. and C.E.N acquired, analysed or interpreted the data. A.S.P., L.H., M.B. and S.H designed research; A.S.P., L.H., S.H. and C.E.N wrote the paper. S.H. and C.E.N supervised the study. All authors have read and approved the submitted version.

## Data availability

The low-coverage genome-wide methylation sequencing data is available at ArrayExpress under accession E-MTAB-14969.

## Supplemental Figures

**Fig. S1. A** Monotherapy dose-response curves for all four AML cell lines following treatment with increasing concentrations of Venetoclax (VEN) for 48 hours or 5-azacytidine (AZA) and Decitabine (DAC) for seven days. Cell viability was assessed by Alamar Blue assay. Mean and standard error of the mean of triplicates for each cell line are shown. **B** Overview of statistical difference between cell lines for the different treatment. P-values were computed using Wilcoxon Rank Sum Tests and adjusted using Benjamini-Hochberg. Significantly adjusted p-values are highlighted in red. **C** Monotherapy dose-response curves for treatment with increasing concentrations of VEN for 24 hours, AZA, DAC, cytarabine (ARA-C), and GSK-3685032 (GSK5032) for three days, and GSK5032 for seven days in all four AML cell lines. Cell viability was determined using Alamar Blue assay. Mean and standard error of the mean from triplicate measurements are shown for each treatment and concentration.

**Fig. S2. A-D** 3D surface plots depicting synergy scores for the combination effects of VEN-AZA, VEN-DAC, VEN-ARA-C, and VEN-GSK5032 across all four AML cell lines. Synergy scores were calculated using the Bliss Independence model. Mean Bliss score and the corresponding p-values are shown above each plot. AZA, 5-azacytidine; ARA-C, Cytarabine; DAC, Decitabine; GSK5032, GSK-3685032; VEN, Venetoclax.

**Fig. S3.** Expression of the methylation-sensitive genes *DAZL* and *GAGE12* was measured by qPCR in KG-1a and MOLM-13 cells after treatment with increasing concentrations of 5-azacytidine (AZA), decitabine (DAC), and GSK-3685032 (GSK5032) for three days. Relative differences in expression compared to the housekeeping gene *GAPDH* are shown.

## Supplemental Data

**Table S1**. Synergy scores for the combination treatments with Venetoclax (VEN) and 5-azacytidine (AZA), decitabine (DAC), cytarabine (ARA-C) for three days or GSK-3685032 (GSK5032) for three and seven days in Kasumi-1, KG-1a, MOLM-13 and THP-1. Synergy scores were calculated using the Bliss Independence model.

**Table S2**. Summary statistics of the synergy scores in Kasumi-1, KG-1a, MOLM-13 and THP-1.

**Table S3**. Sequencing metrics for low pass whole methylome sequencing of Kasumi-1, KG-1a, MOLM-13 and THP-1 after treatment with increasing concentrations of 5-azacytidine (AZA), decitabine (DAC), cytarabine (ARA-C) for three days or GSK-3685032 (GSK5032) for three and seven days.

**Table S4**. Average CpG methylation as percentage, globally and at LINE elements, assessed by low pass whole methylome sequencing of Kasumi-1, KG-1a, MOLM-13 and THP-1 after treatment with increasing concentrations of 5-azacytidine (AZA), decitabine (DAC), cytarabine (ARA-C) for three days or GSK-3685032 (GSK5032) for three and seven days.

**Table S5**. Average coverage per CpG, assessed by low pass whole methylome sequencing of Kasumi-1, KG-1a, MOLM-13 and THP-1 after treatment with increasing concentrations of 5-azacytidine (AZA), decitabine (DAC), cytarabine (ARA-C) for three days or GSK-3685032 (GSK5032) for three and seven days.

**Table S6**. Average mean viability and global CpG methylation for Kasumi-1, KG-1a, MOLM-13 and THP-1 after treatment with increasing concentrations of 5-azacytidine (AZA), decitabine (DAC), cytarabine (ARA-C) for three days or GSK-3685032 (GSK5032) for three and seven days.

