## Supplemental Figures 1-3 for "DNA Demethylation Is Dispensable for Venetoclax-HMA Synergy in Acute Myeloid Leukaemia"

### Supplemental Figure 1

A

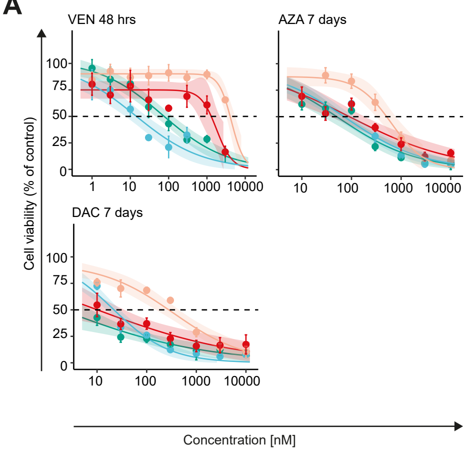


C

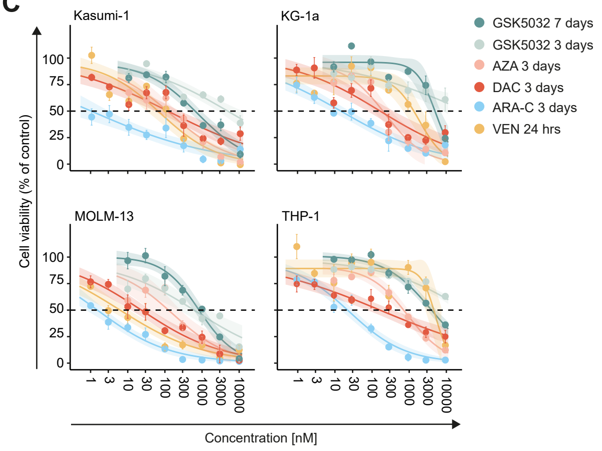


B

| | VEN 24 hrs | VEN 48 hrs | ARA-C 3 days | AZA 3 days | AZA 7 days | DAC 3 days | DAC 7 days | GSK5032 3 days | GSK5032 7 days |
|---|---|---|---|---|---|---|---|---|---|
| Kasumi-1 *vs* KG-1a | 0.096 | 0.476 | 0.117 | 0.117 | 0.059 | 0.117 | 0.059 | 0.117 | 0.059 |
| Kasumi-1 *vs* MOLM-13 | 0.240 | 0.147 | 0.062 | 0.062 | 0.754 | 0.062 | 0.754 | 0.062 | 0.075 |
| Kasumi-1 *vs* THP-1 | **0.003** | **0.015** | 0.062 | 0.062 | **0.003** | 0.062 | 0.003 | 0.062 | **0.003** |
| KG-1a *vs* MOLM-13 | **0.003** | 0.051 | **0.001** | **0.001** | 0.129 | **0.001** | 0.129 | **0.001** | 0.129 |
| KG-1a *vs* THP-1 | 0.096 | **0.014** | 0.670 | 0.670 | 0.226 | 0.670 | 0.225 | 0.670 | 0.226 |
| MOLM-13 *vs* THP-1 | **1.467e-05** | **0.0003** | **0.0004** | **0.0004** | **0.014** | **0.0004** | **0.014** | **0.0004** | **0.014** |

### Supplemental Figure 2

**A**

**Kasumi-1**

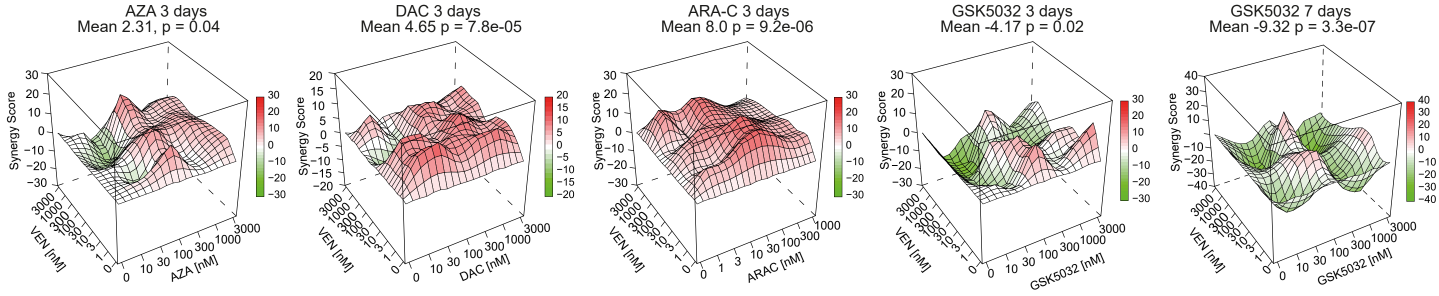


**B**

**KG-1a**

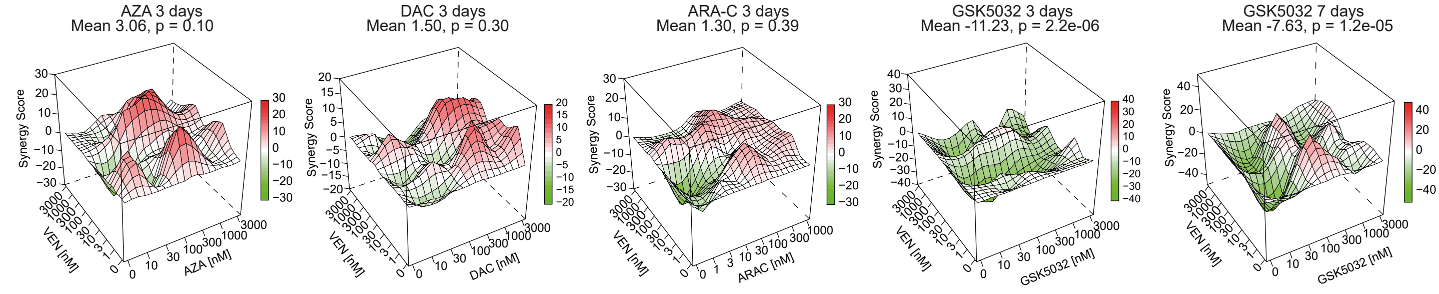


**C**

**MOLM-13**

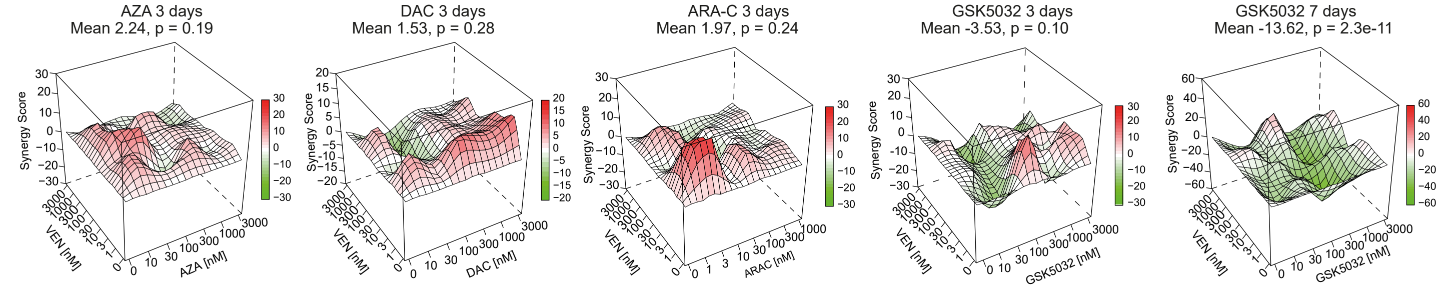


**D**

**THP-1**

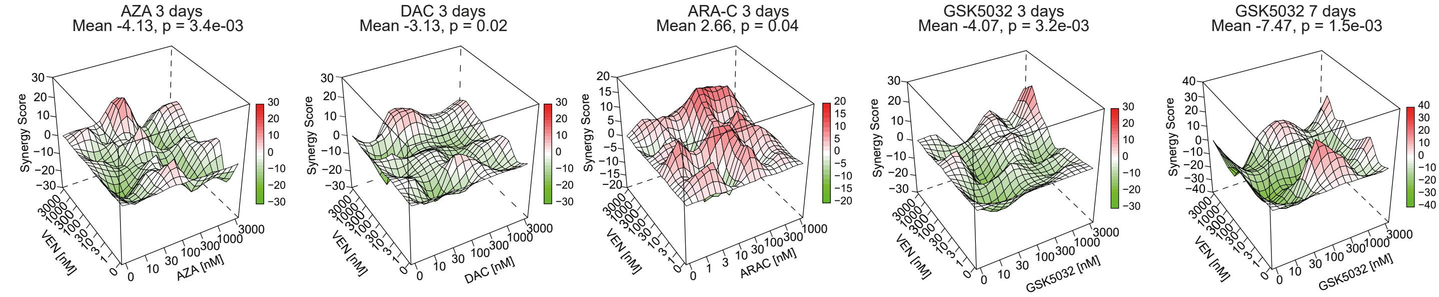

### Supplemental Figure 3

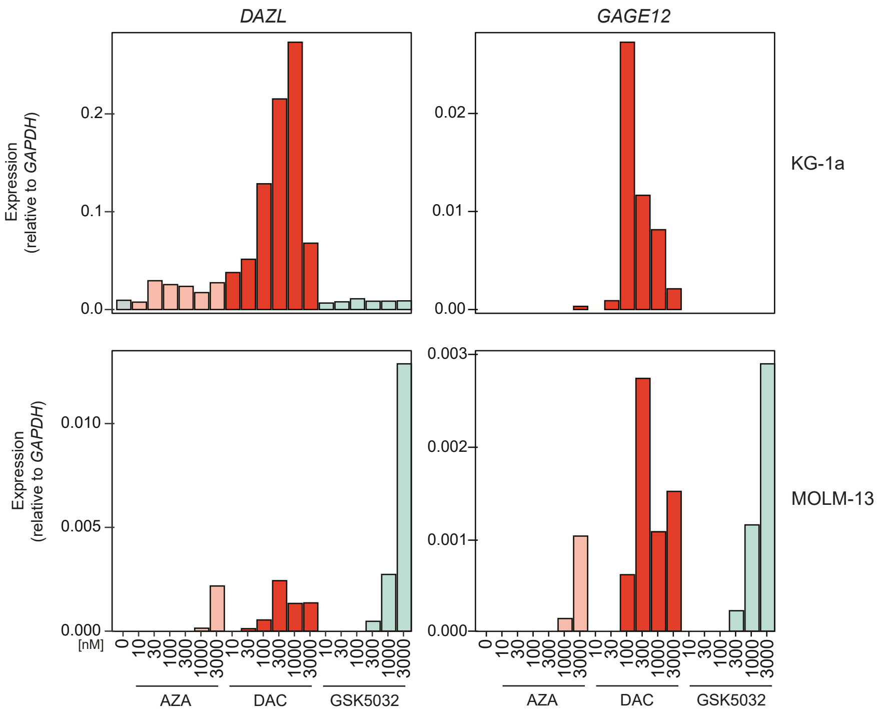
